# Co-contraction index improves estimates of knee joint contact forces: A musculoskeletal modelling study

**DOI:** 10.64898/2026.05.28.728089

**Authors:** Sabrina Hörmann, Nazli Tümer, Amir A. Zadpoor, Ajay Seth

**Affiliations:** Delft University of Technology, Department of Biomechanical Engineering, Delft, The Netherlands

## Abstract

Healthy individuals are hypothesized to adopt muscle coordination strategies that minimize energy expenditure. However, under pathological conditions, these patterns often change, as patients adopt alternative strategies to enhance stability. In musculoskeletal modeling, such changes in muscle coordination are often not accounted for when estimating internal quantities such as knee joint contact forces. To address this, we adapted the objective function of a muscle redundancy solver to inform muscle activations from electromyography measurements, minimizing errors in both muscle activations and co-contraction levels. The resulting estimates of knee joint contact force were compared with *in vivo* measurements across three activities. The co-contraction index-informed approach achieved the lowest root-mean-square-error of 0.31 body weight, averaged across all subjects and activities, improving the results of the minimum activation solver by 7% body weight. Notably, root-mean-squared-error increased with the level of co-contraction with the minimum activation approach (*β* = 1.55, 95% confidence interval [0.79, 2.30]), whereas the co-contraction index-informed approach remained less sensitive (*β* = 0.78, 95% confidence interval [0.14, 1.43]). These findings suggest that objective functions based on minimum muscle activation may be insufficient to accurately estimate knee joint forces in individuals with altered muscle coordination. Incorporating co-contraction information is therefore essential to capture subject-specific adaptations.

## Introduction

Non-pharmacological rehabilitation programs for patients with knee osteoarthritis (KOA) primarily focus on therapeutic exercises to increase muscle strength, improve gait function, and enhance balance^1^. Currently, these exercise recommendations are largely based on expert opinion, and there is only limited evidence supporting specific movement programs^2^. Therefore, one question that often arises for both physicians and patients is the appropriate amount and intensity of exercise that is safe and beneficial for the patient^3^. While exercise interventions have been shown to reduce pain and improve both functional outcomes and quality of life compared to no intervention^4,5^, MRI findings indicate no corresponding reduction in joint inflammation^4^. Excessive knee joint loading is closely associated with cartilage inflammation, which is thought to be a factor in the progression of KOA^6–8^. This suggests that conventional training protocols are not sufficiently focused on reducing knee joint loading during daily activities.

Gait retraining has been reported as an effective approach to redistributing or reducing knee joint forces^9–11^. Strategies for adapting walking patterns described in the literature include altering the foot progression angle^10,12^, using a medial thrust gait^13^, or using trunk lean^13^. Furthermore, adapting muscle coordination strategies, such as activating the soleus rather than the gastrocnemius during the push-off phase of walking, has been reported to decrease knee joint contact forces at the second peak^9^. These studies, therefore, indicate how essential it is to track the subject’s knee joint loading to evaluate the effect of using gait retraining during rehabilitation and to assess the progression of KOA. Since direct measurements of knee joint forces in an intact knee are not feasible, musculoskeletal (MSK) modeling is a viable approach for estimating knee joint contact forces during gait retraining.

One key challenge in MSK modeling is solving the muscle redundancy problem. This problem arises because the human musculoskeletal system has more muscles than the number of degrees of freedom required to produce a specific movement. Consequently, different muscle activation strategies can produce the same motion but result in varying internal quantities, such as joint loading and metabolic cost, posing a challenge for determining the strategy an individual subject adopts. Healthy adults are thought to select a neuromuscular activation strategy that minimizes metabolic cost during tasks such as walking^14^. This strategy is implemented by computing the muscle activation pattern that minimizes the sum of squared muscle activations^15^. However, KOA patients may optimize muscle coordination strategies, for example, to reduce pain or enhance joint stability, thereby reducing the risk of falls or trips^16^. This adaptation in muscle coordination strategy is supported by previous findings indicating greater muscle co-contraction in some individuals with early-stage KOA than in healthy controls^17,18^. Increases in muscle co-contraction, commonly quantified using a co-contraction index (CCI), elevates knee joint loading and should therefore be considered in knee joint contact force estimations of KOA patients^19,20^. To account for differences in the muscle coordination strategy, the simulation framework must be informed of the patient-specific differences in the muscle activation pattern. The most effective non-invasive approach to measure patient-specific muscle activation patterns is electromyography (EMG). EMG-informed simulations integrate neuromuscular control strategies by utilizing surface EMG signals recorded from selected muscles, thereby providing subject-specific insight into muscle activation dynamics. The application of current EMG-informed approaches in out-of-laboratory settings remains limited, primarily due to the requirement for EMG data from ideally all relevant muscles^21^ or, at a minimum, from eight muscles^22^. In addition, these approaches typically require subject-specific model calibration, which can be computationally demanding^23,24^.

In this study, we explore an alternative approach by incorporating EMG measurements directly into the objective function of the muscle redundancy solver^9^, aiming to reduce the need for extensive EMG recordings and time-consuming calibration procedures. The primary objective of this study is to test estimated knee contact forces (KCF) from MSK modeling during level walking, ramp descent, and stair descent, against *in vivo* measurements from instrumented implants^25,26^.

## Methods

To achieve this, we used the measurements from the publicly available Comprehensive Assessment of the Musculoskeletal System (CAMS) dataset^26^ as inputs to our MSK modeling simulation pipeline to estimate KCF. The resulting KCF estimates obtained from minimum muscle activation, EMG-informed, and CCI-informed solvers were compared to the *in vivo* data and then quantitatively evaluated. An overview of the methods is presented in Figure 1.

**Figure 1.**
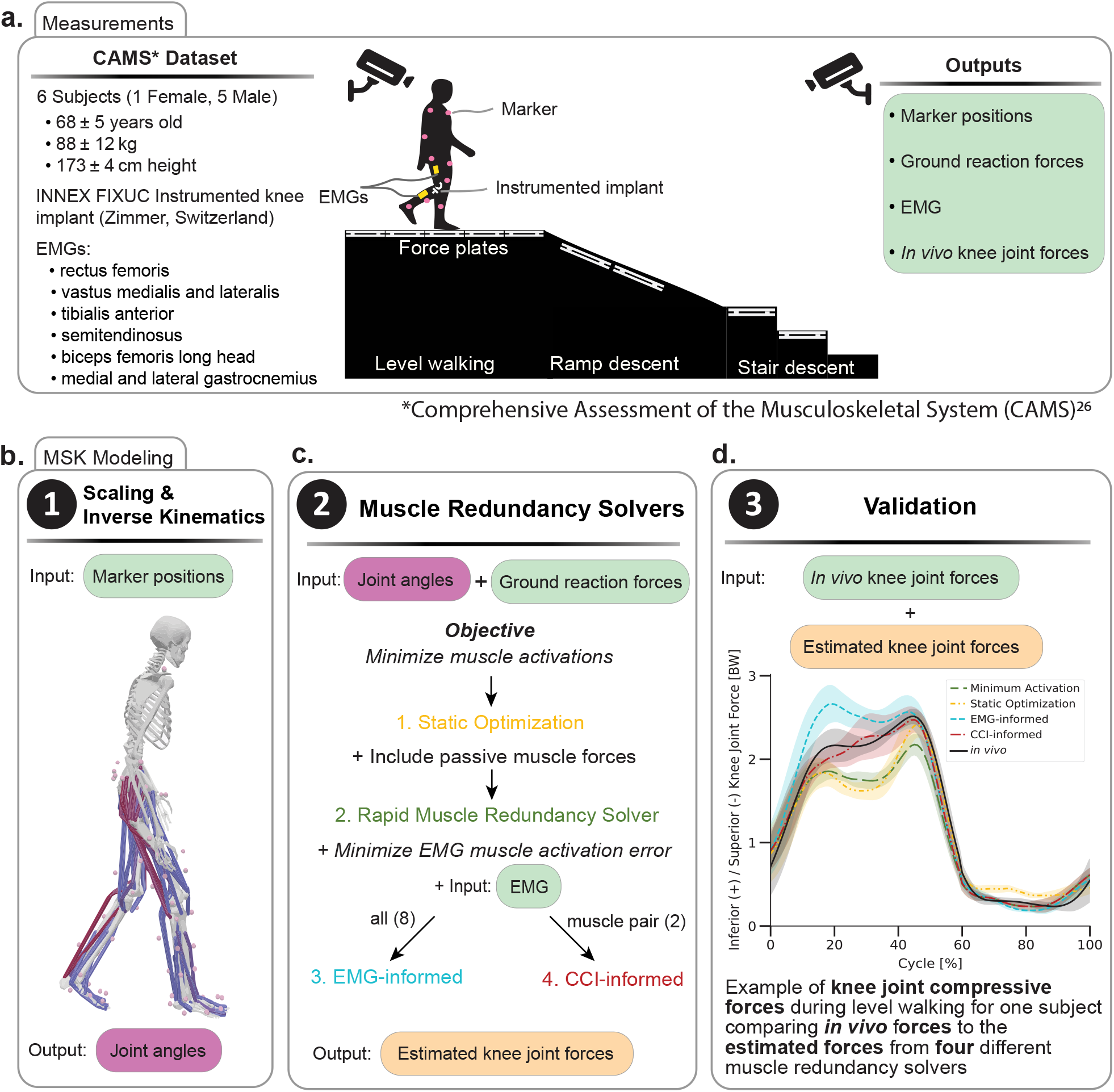
Overview of the experimental and simulation workflow. (**a**) Input measurements are obtained from the publicly available CAMS dataset^26^. All measurements from the CAMS dataset are marked in green. (**b**) Marker trajectories are used to scale a generic musculoskeletal model and perform inverse kinematics. (**c**) The resulting joint angles, together with measured ground reaction forces, are used as input for four muscle redundancy solvers. Static Optimization and the Rapid Muscle Redundancy solver minimize muscle activation; the latter additionally accounts for passive muscle forces. For the EMG-informed and CCI-informed solvers, the objective function additionally minimizes the error between predicted and measured muscle activations. The EMG-informed solver uses measurements from eight muscles as input, whereas the CCI-informed solver minimizes the error in co-contraction of Medial Gastrocnemius and Vastus Lateralis. (**d**) All solvers estimate knee joint compressive forces, which are subsequently tested against *in vivo* measurements using an instrumented implant.

### Experimental Data

The CAMS knee dataset^26^ provides *in vivo* measurements of the three force and three moment components (90–100 Hz acquisition frequency) acting on the tibial implant. The dataset includes measurements of five males and one female (68 ± 5 years old, 88 ± 12 kg, and 173 ± 4 cm height), who received cemented INNEX knee implants (Zimmer, Switzerland; FIXUC). We analyzed all gait-related activities including level walking, ramp descent, and stair descent. For level walking, simulations were conducted on eight trials per subject. For ramp and stair descent, four trials per subject were analyzed. Data from subject K3R during ramp descent were excluded due to invalid ground reaction force measurements, as both feet contacted a single force plate during these trials. Whole-body kinematics were measured using 75 skin markers and a 26-camera motion-capture system (Vicon, UMG, UK) at 100 Hz. Ground reaction forces were collected at 2000 Hz with six force plates embedded in the walkway (Kistler Instrumentation, Winterthur, Switzerland). Bilateral muscle activity for eight major lower limb muscles (rectus femoris, vastus medialis, vastus lateralis, tibialis anterior, semitendinosus, biceps femoris long head, medial gastrocnemius, and lateral gastrocnemius) was detected using a 16-channel wireless EMG system (Trigno, Delsys, USA) with a signal delay of 48ms. The raw EMG was high-pass filtered at 100 Hz, full-wave rectified, and low-pass filtered at 4 Hz to obtain EMG envelopes, as per ISEK^27^. The 48 ms signal delay was corrected. Furthermore, the ground reaction forces were low-pass filtered (zero-phase shift Butterworth filter, 4th order, 12Hz).

### Musculoskeletal Modeling

We used a generic full-body MSK model described by Rajagopal et al.^28^, with adaptations from Lai et al.^29^ and Uhlrich et al.^9^, incorporating 37 degrees of freedom (DOF), 80 musculotendon actuators for the lower extremities, and 17 ideal torque actuators for the upper body. The model included six DOF between the pelvis and the ground, three rotational DOF between the pelvis and torso, three rotational DOF at each hip, one rotational DOF at each knee that parametrized the remaining rotational and translational DOF of the tibiofemoral and patellofemoral joints with respect to knee flexion, and one rotational DOF at each of the ankle, subtalar, and metatarsophalangeal joints.

MSK modeling and simulation were performed using OpenSim 4.5^30,31^, with the simulation workflow automated using the Snakemake workflow management system^32^. The generic model was scaled using the OpenSim Scale Tool to match subject-specific anthropometric measurements obtained from a standing static trial, during which virtual model markers were adjusted to align with experimental marker locations. Model scaling proportionally adjusted muscle optimal fiber lengths, tendon slack lengths, and muscle moment arms according to segment-specific scaling factors, while preserving the ratio of tendon slack length to optimal fiber length for each muscle. Model kinematics were estimated from experimental marker data using the OpenSim Inverse Kinematics tool, which minimizes the distance error between experimental marker trajectories and corresponding virtual model markers. The resulting joint kinematics were used as inputs to the static optimization (SO) solver and the Rapid Muscle Redundancy (RMR)^33^ solver to estimate muscle activations. Knee joint contact forces were computed from muscle force estimates using the OpenSim Joint Reaction Analysis tool. For muscle forces estimated with the RMR solver, the corresponding knee joint contact forces were computed in Python (version 3.10.17) by calling the OpenSim API. To ensure consistency with the orientation of the *in vivo* implant, the reference systems were adjusted according to the reported patient-specific tibial slope^26^.

### Muscle redundancy solvers

To capture the effect of changes in muscle coordination patterns in KOA patients, we adapted the objective function of the RMR solver to account for the measured EMG signals. The SO and the RMR solver both address the muscle redundancy problem by identifying the muscle activation *a*_*i,t*_, where *i* ∈{1,&, *N*_*m*_} refers to each muscle, that minimizes the cost function. The cost function is defined as the sum of squared muscle activations across all muscles *N*_*m*_ at each time step *t* of the input motion (Equation 1–SO and Equation 2–RMR)^15,33,34^.

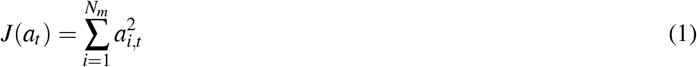

The optimization is subject to constraints ensuring that the model dynamics (equations of motion) produce accelerations consistent with the experimentally measured motion. In addition, muscle-generated joint moments were limited to match those computed from inverse dynamics, and muscle forces were kept within physiological boundaries defined by the maximal isometric muscle force. Furthermore, for each DOF in the MSK model, ideal reserve actuators were included to compensate for residual dynamics resulting from inaccuracies and inconsistencies between the model and the experimental data. For SO, limiting the activation of the reserve actuators is achieved by defining a small optimal force (*<* 10 Nm), whereas in the RMR solver, the reserve actuators are separately included in the objective function (Equation 2). To discourage the solver from using the reserve actuator *r* _*j,t*_, where *j* ∈ {1,&, *N*_*n*_} corresponds to each coordinate actuator, the reserve actuator weights *v*_*j*_ are set proportionally higher, at 10 times the muscle weights *w*_*i*_.

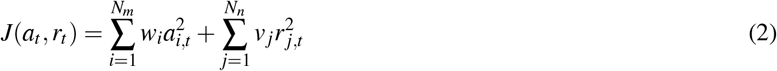

Furthermore, unlike the SO solver, the RMR solver incorporates passive muscle fiber forces to preserve physiological behaviors and imposes constraints on muscle activation dynamics to ensure continuity of the activation profile^33^.

In the next step, we extended the RMR solver’s cost function to account for tracking EMG measurements. Therefore, we included a third term in the cost function that minimizes the error between the normalized EMG measurement and the estimated muscle activation (Equation 3). The optimization problem was formulated as

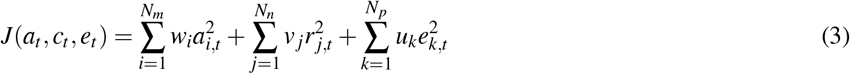

where *e*_*k,t*_ = *a*_*k,t*_ − EMG_norm*k,t*_ and *k* ∈ {1,&, *N*_*p*_} corresponds to the muscles for which EMG measurements are available.

To penalize deviations between estimated muscle activations *a*_*k,t*_ and EMG signals normalized to maximal voluntary contraction (MVC) EMG_norm*k,t*_, the weight *u*_*k*_ were scaled to be twice as large as the muscle weights *w*_*i*_. Thus, fidelity to measured activation patterns is prioritized over minimization of muscle activations. As EMG data were available only for the rectus femoris, vastus medialis, vastus lateralis, semitendinosus, biceps femoris long head, medial gastrocnemius, lateral gastrocnemius, and tibialis anterior, the EMG measurement-to-simulation estimate error term was evaluated exclusively for these muscles. For all remaining muscles, no additional EMG cost term was included.

Finally, we introduce a CCI-informed cost designed to quantify the level of co-contraction contributing to elevated KCF. The CCI is a relative measure and is therefore less sensitive to inaccuracies in MVC normalization that arise when subjects are unable to achieve true maximal muscle contractions. As a result, no additional calibration is required to correct for MVC-related bias, avoiding the need for further measurements. The initial RMR cost function (Equation 1) was then extended by an additional term that minimizes the squared error between the CCI calculated from EMG measurements CCI_measured,*t*_ and from the simulation CCI_simulation,*t*_ (Equation 4). The objective function was defined as

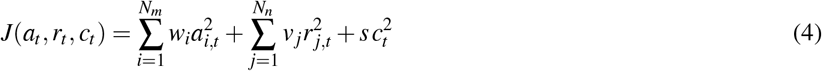

where *c*_*t*_ = CCI_simulation,*t*_ − CCI_measured,*t*_.

Similar to the EMG-informed approach, the weighting factor *s* for the CCI error term was set to twice the muscle weights *w*_*i*_, thereby prioritizing minimization of CCI errors over minimization of muscle activation to account for changes in muscle coordination strategy.

The CCI was calculated as proposed by Rudolph et al.^19^:

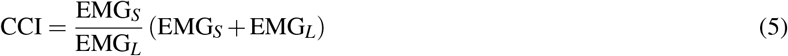

where EMG_*S*_ denotes the smaller muscle activity value and EMG_*L*_ the larger muscle activity value, which prevents division by zero and constrains the CCI to the range 0–2^19^. The CCI was calculated for multiple antagonistic muscle pairs crossing the knee joint. The largest discrepancy between measured and RMR predicted co-contraction based on the minimum activation objective function was observed between the vastus lateralis (VL) and medial gastrocnemius (MG) pair. Furthermore, both muscles are superficially located, which enhances the reliability of surface electromyography measurements due to reduced signal attenuation from overlying tissues^35^. Therefore, this muscle pair was selected for use in the CCI-informed simulations.

### Statistical Analysis and Evaluation Metrics

Three statistical measures were utilized to evaluate the correspondence between the simulated and *in vivo* KCF. The root mean squared error (RMSE) quantifies the overall magnitude of the error between the simulated and measured forces across all time steps within each trial. The coefficient of determination R^2^ describes how well the temporal waveform of the estimated forces corresponds to the *in vivo* measurements. In addition, the peak force error was defined as the difference at the second peak of the stance phase, computed as the maximum absolute value within 40–60% of the trial for both the measured and simulated knee joint compressive forces. For each metric, we calculated the mean and standard deviation over all trials, including level walking, ramp descent, and stair descent. Statistical differences were assessed using the Wilcoxon signed-rank test for paired, non-normally distributed data, with Holm–Bonferroni correction applied to account for multiple comparisons across muscle redundancy solvers. A corrected *p*-value below *α* = 0.05 was considered statistically significant. Additionally, we analyzed the relationship between RMSE and CCI for the four muscle redundancy solvers. First, a linear mixed-effects regression model was specified with a subject-specific random intercept to account for between-subject variability in baseline RMSE across repeated trials. However, the model did not converge consistently across analyses and therefore did not yield reliable estimates for inference. As a result, we evaluated the relationship between RMSE and CCI using linear regression with cluster-robust standard errors to account for within-subject dependence in the data. The six subjects were treated as clusters. From this analysis, we report the slope (*β*, in BW per unit CCI), cluster-robust standard errors, 95 % confidence intervals (95 % CI), *p*-values, and the number of observations (*N*). Statistical significance was defined as *p <* 0.05. All statistical analyses were performed in Python (version 3.10.17) using the SciPy library (version 1.15.2). During post-processing, the compressive KCF was smoothed using a Savitzky-Golay filter (window length = 41 frames, polynomial order = 3) to reduce numerical artifacts in the estimated KCF.

## Results

In this study, we compared the estimated compressive KCF obtained from four muscle redundancy solvers - SO, minimum activation (RMR), EMG-informed, and CCI-informed – to the *in vivo* measured KCF from all six subjects in the CAMS dataset. The comparison was performed across three activities: level walking, ramp descent, and stair descent. To ensure consistency across simulations, a fixed weighting scheme was applied in the objective function of the RMR solver. Specifically, the weight of the muscle activation minimization term was used as the baseline reference and set to 1, and all other cost function terms were scaled accordingly. Therefore, the reserve actuator term was assigned a weight of 10, while the EMG-informed and CCI-informed cost function terms were assigned each a weighting factor of 2.

The resulting compressive KCF profiles are shown in Figure 2, illustrating the mean ± 1 standard deviation (std) across multiple trials over time for each subject during level walking (Figure 2a), ramp descent (Figure 2b), and stair descent (Figure 2c). Overall, all four muscle redundancy solvers captured the general shape and trends of the *in vivo* measurements. To further quantify differences between solvers, evaluation metrics were calculated against the ground truth from the instrumented implants.

**Figure 2.**
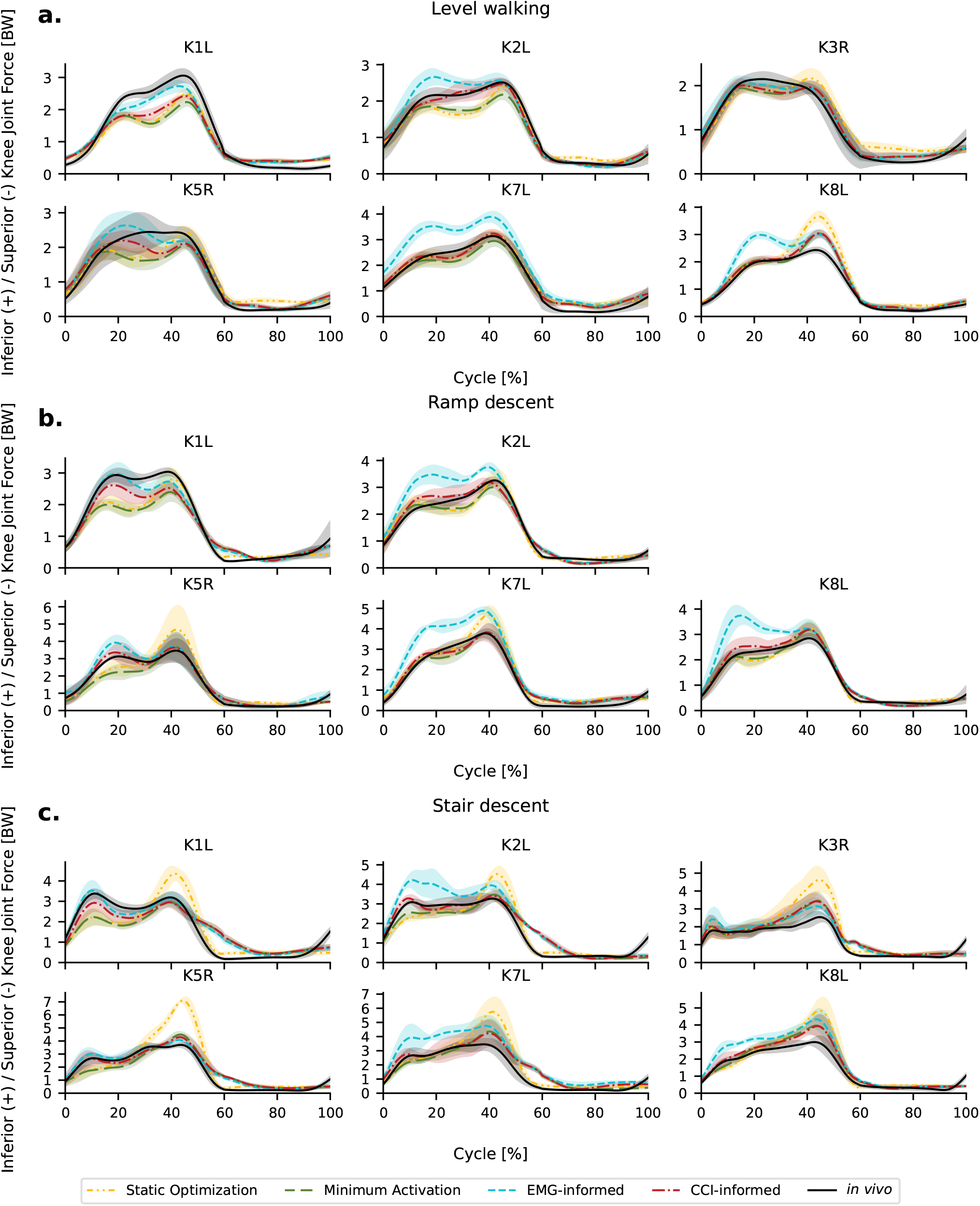
Simulated and measured compressive knee joint contact forces over the activity cycle. Comparison of estimated knee contact forces (mean ± 1 std) from Minimum Activation (green), Static optimization (yellow), EMG-informed (blue) and CCI-informed (red) to *in vivo* measurements (black) from an instrumented implant for each subject in level walking (**a**), ramp descent (**b**) and stair descent (**c**).

The resulting analysis is summarized in Figure 3a, which presents the RMSE across all included trials and activities for each subject, as well as the overall across all subjects. In addition, Table 1 reports the corresponding mean ± std RMSE values. The results indicate that CCI-informed simulations achieved the lowest overall RMSE, with a mean value of 0.31 body weight (BW). For individual subjects, the CCI-informed approach showed the lowest RMSE for K2L, K7L, and K8L. For the remaining subjects, K1L, K3R, and K5R, the EMG-informed approach achieved the lowest RMSE. However, the EMG-informed solver showed the highest RMSE for K2L, K7L, and K8L, resulting in an overall mean RMSE of 0.44 BW. The minimum activation (RMR) simulations resulted in the second-lowest overall RMSE (0.38 BW), 0.07 BW above the CCI-informed simulations. Comparison between CCI-informed and minimum activation (RMR) approach revealed significant improvements (*p <* 0.05) in RMSE using the CCI-informed approach overall and for subjects K1L, K2L, K5R, and K7L, whereas the other two subjects showed only minimal, non-significant reductions. The largest improvements compared to the minimum activation (RMR) approach were observed in subject K1L and K5R, with a descrease in mean RMSE of 0.14 BW. Across all subjects combined, static optimization exhibited the highest mean RMSE of 0.47 BW.

**Table 1.**
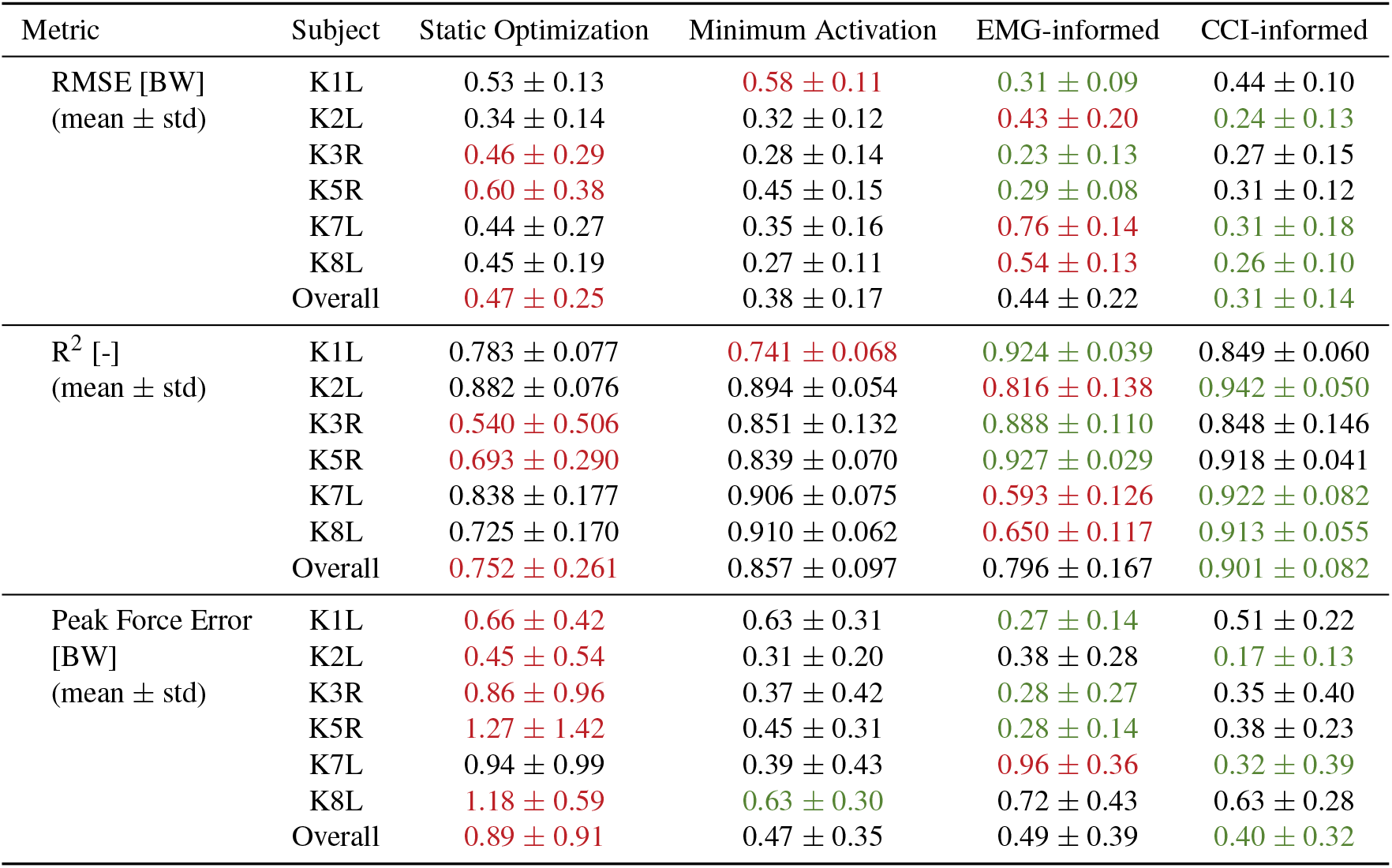
Summary of RMSE, *R*^2^ and peak force error of the second peak in stance phase for each subject and across all subjects (Overall). Metrics are compared across four muscle redundancy solvers: Static Optimization, Minimum Activation (RMR), EMG-informed, and CCI-informed. For each subject and the overall results, the best-performing solver is highlighted in green and the worst-performing solver in red. The results show that the CCI-informed simulations performed best for each evaluation metric for all subjects combined (Overall).

| Metric | Subject | Static Optimization | Minimum Activation | EMG-informed | CCI-informed |
| --- | --- | --- | --- | --- | --- |
| RMSE [BW]<br>(mean ± std) | K1L | 0.53 ± 0.13 | 0.58 ± 0.11 | 0.31 ± 0.09 | 0.44 ± 0.10 |
| | K2L | 0.34 ± 0.14 | 0.32 ± 0.12 | 0.43 ± 0.20 | 0.24 ± 0.13 |
| | K3R | 0.46 ± 0.29 | 0.28 ± 0.14 | 0.23 ± 0.13 | 0.27 ± 0.15 |
| | K5R | 0.60 ± 0.38 | 0.45 ± 0.15 | 0.29 ± 0.08 | 0.31 ± 0.12 |
| | K7L | 0.44 ± 0.27 | 0.35 ± 0.16 | 0.76 ± 0.14 | 0.31 ± 0.18 |
| | K8L | 0.45 ± 0.19 | 0.27 ± 0.11 | 0.54 ± 0.13 | 0.26 ± 0.10 |
| | Overall | 0.47 ± 0.25 | 0.38 ± 0.17 | 0.44 ± 0.22 | 0.31 ± 0.14 |
| $R^2$ [-]<br>(mean ± std) | K1L | 0.783 ± 0.077 | 0.741 ± 0.068 | 0.924 ± 0.039 | 0.849 ± 0.060 |
| | K2L | 0.882 ± 0.076 | 0.894 ± 0.054 | 0.816 ± 0.138 | 0.942 ± 0.050 |
| | K3R | 0.540 ± 0.506 | 0.851 ± 0.132 | 0.888 ± 0.110 | 0.848 ± 0.146 |
| | K5R | 0.693 ± 0.290 | 0.839 ± 0.070 | 0.927 ± 0.029 | 0.918 ± 0.041 |
| | K7L | 0.838 ± 0.177 | 0.906 ± 0.075 | 0.593 ± 0.126 | 0.922 ± 0.082 |
| | K8L | 0.725 ± 0.170 | 0.910 ± 0.062 | 0.650 ± 0.117 | 0.913 ± 0.055 |
| | Overall | 0.752 ± 0.261 | 0.857 ± 0.097 | 0.796 ± 0.167 | 0.901 ± 0.082 |
| Peak Force Error<br>[BW]<br>(mean ± std) | K1L | 0.66 ± 0.42 | 0.63 ± 0.31 | 0.27 ± 0.14 | 0.51 ± 0.22 |
| | K2L | 0.45 ± 0.54 | 0.31 ± 0.20 | 0.38 ± 0.28 | 0.17 ± 0.13 |
| | K3R | 0.86 ± 0.96 | 0.37 ± 0.42 | 0.28 ± 0.27 | 0.35 ± 0.40 |
| | K5R | 1.27 ± 1.42 | 0.45 ± 0.31 | 0.28 ± 0.14 | 0.38 ± 0.23 |
| | K7L | 0.94 ± 0.99 | 0.39 ± 0.43 | 0.96 ± 0.36 | 0.32 ± 0.39 |
| | K8L | 1.18 ± 0.59 | 0.63 ± 0.30 | 0.72 ± 0.43 | 0.63 ± 0.28 |
| | Overall | 0.89 ± 0.91 | 0.47 ± 0.35 | 0.49 ± 0.39 | 0.40 ± 0.32 |

**Figure 3.**
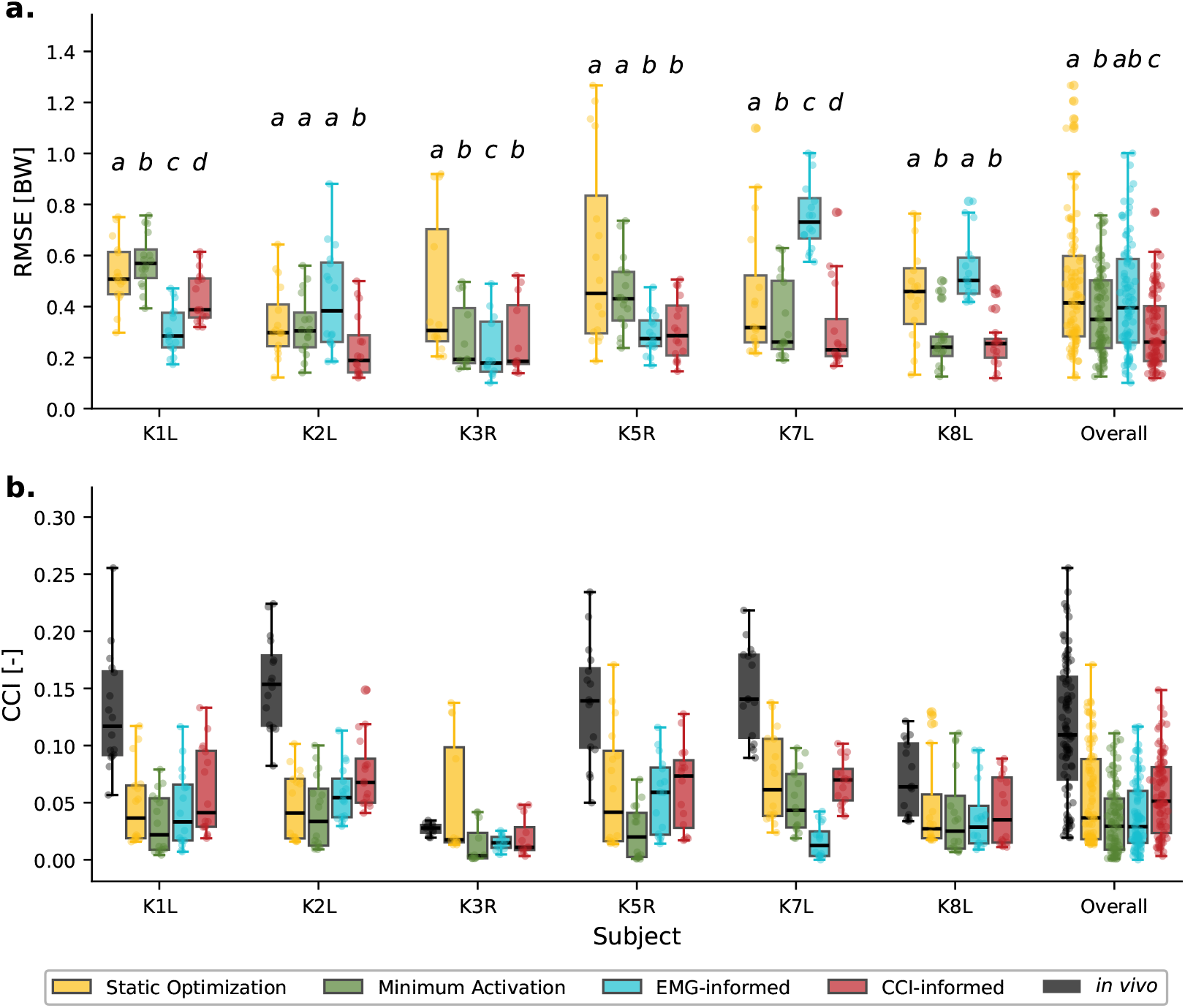
Comparison of RMSE and CCI across muscle redundancy solvers for each subject and across all subjects combined (Overall) including all three activities. (**a**) RMSE of estimated compressive KCF from four muscle redundancy solvers across all activities, shown per subject and all subjects combined (Overall). Boxplots represent the distribution of RMSE values for each subject across all trials from level walking, ramp descent and stair descent, as well as the overall distribution across subjects. Letters above each boxplot indicate significant differences *p <* 0.05 in RMSE within each grouped boxplot. Different letters denote a significant difference, while identical letters indicate no significant difference. *P*-values are reported in Table S1 of the Supplementary Material. CCI-informed (red) simulations result in the lowest overall RMSE. The minimum activation (green) solver yields mean RMSE values between 0.27 and 0.58 BW. EMG-informed (blue) simulations results in considerable inter-subject variability: RMSE is lowest for K1L, K3R, and K5R, but highest for K2L, K7L, and K8L. Static Optimization (yellow) simulations lead to the highest RMSE across solvers. (**b**) CCI calculated from Vastus Lateralis and Medial Gastrocnemius muscle activations, comparing EMG measurements (black) with co-contraction predicted by four muscle redundancy solvers across all activities. Boxplots are shown per subject and for the overall distribution across subjects. While *in vivo* CCI is low for K3R and K8L, co-contraction in the other subjects is considerably higher.

In addition, the corresponding CCI values of the VL and MG across all trials and activities for each subject, as well as the overall across subjects are summarized in Figure 3b. *In vivo* EMG measurements indicate higher co-contraction for subjects K1L, K2L, K5R, and K7L compared to the estimated co-contraction obtained with the minimum activation (RMR) approach. In CCI-informed simulations, this discrepancy is reduced by an average of 0.03 in these subjects. For the remaining subjects, the measured and estimated co-contraction values are more closely aligned across all muscle redundancy solvers.

Furthermore, Table 1 reports mean ± std RMSE, *R*^2^, and peak force error at the second peak of the stance phase across activities for each subject individually, as well as the overall across subjects. The *R*^2^ results show that the CCI-informed simulations achieve an overall value above 0.9, which is also reflected in the closer agreement with the *in vivo* measurements, for example in the first peak during ramp descent and stair descent for subject K1L, as shown in Figures 2b and 2c. In contrast, each of the other four muscle redundancy solvers yielded an overall *R*^2^ below 0.86. For the peak force error, similar patterns to those observed for the RMSE can be observed between the muscle solvers. An exception is the SO solver, which exhibits the highest peak force error in five of the six subjects, which is also reflected in the pronounced peak deviations in the stair descent curves shown in Figure 2c. The overall peak error of SO is 0.89 BW, which is 0.49 BW higher compared to the best-performing CCI-informed simulations, which show an overall error of 0.4 BW.

In the next step, we analyzed the RMSE between the *in vivo* measured KCF and the estimates from the four muscle redundancy solvers across all trials and subjects for each activity, as presented in Figure 4a. In addition, Table S2 of the Supplementary Material reports the corresponding mean ± 1 std RMSE values. For the SO, minimum activation (RMR), and CCI-informed solvers, RMSE show minimal change between level walking and ramp descent, whereas all three solvers show an increase in mean RMSE during stair descent compared to the other two activities. For the EMG-informed solver, mean RMSE progressively increases from level walking to ramp descent and stair descent. Moreover, CCI-informed simulations differed significantly from the other three solvers (*p <* 0.01), exhibiting a 0.07 BW reduction compared to minimum activation (RMR) approach in mean RMSE for each activity.

**Figure 4.**
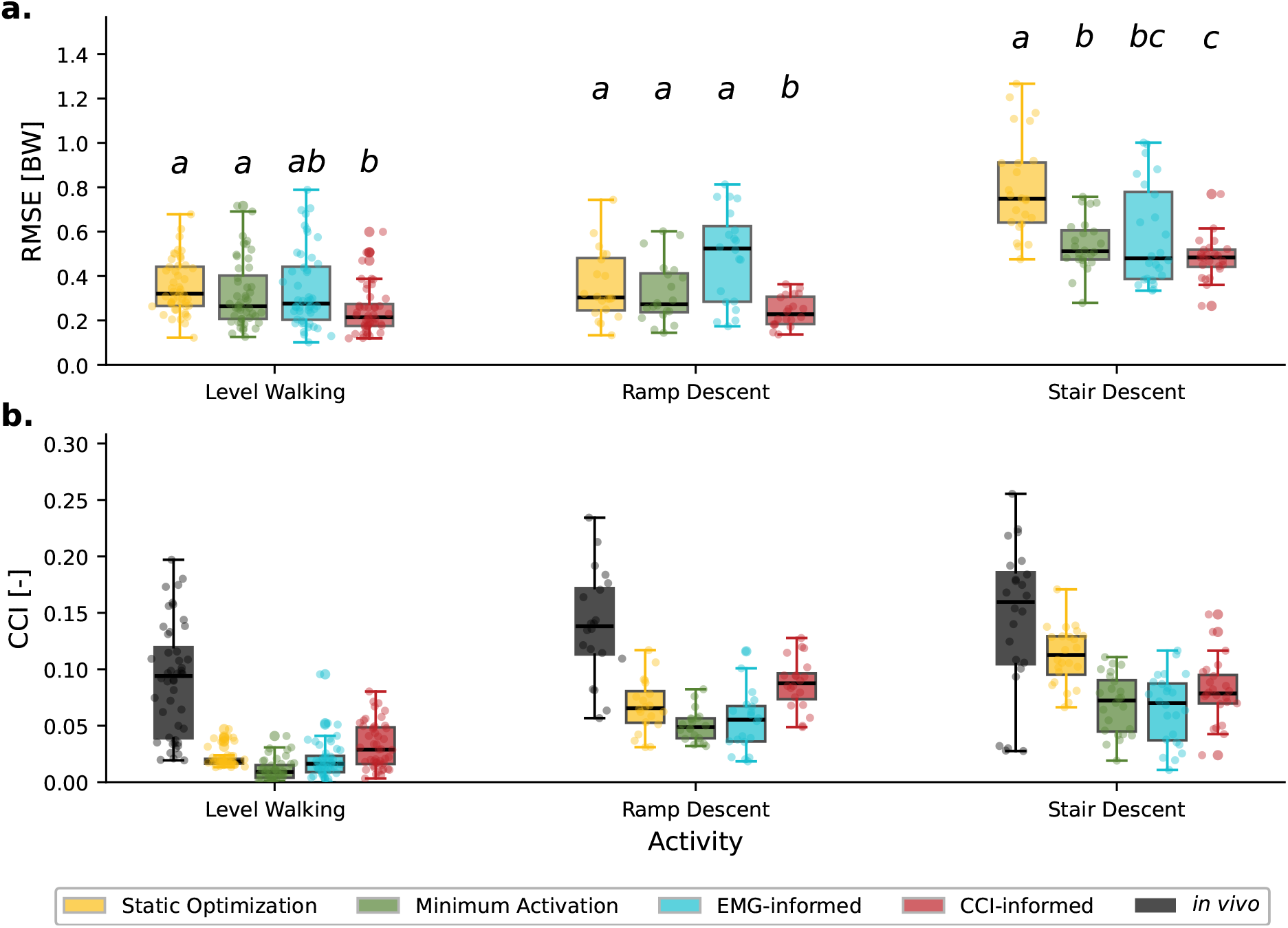
Comparison of RMSE and CCI across muscle redundancy solvers for each activity including all six subjects. **(a)**RMSE of estimated compressive KCF from four muscle redundancy solvers across all subjects, shown per activity. Boxplots represent the distribution of RMSE values for each activity (level walking, ramp descent, stair descent) across all trials from six subjects. Letters above each boxplot indicate significant differences (*p <* 0.05) in RMSE within each grouped boxplot. Different letters denote a significant difference, while identical letters indicate no significant difference. *P*-values are reported in Table S3 of the Supplementary Material. For Static Optimization (yellow), Minimum Activation (green), and CCI-informed (red), RMSE is similar between level walking and ramp descent but increases during stair descent. EMG-informed (blue) simulations show a progressive increase in RMSE from level walking to ramp descent and further to stair descent. **(b)** Vastus Lateralis and Medial Gastrocnemius CCI comparing EMG measurements (black) to resulting co-contraction from four different muscle redundancy solvers across all subjects, shown per activity. *In vivo* CCI results show increasing co-contraction from level walking to ramp descent and further to stair descent.

Furthermore, Figure 4b presents the corresponding CCI values of the VL and MG across all subjects and trials for each activity. The results show that *in vivo* CCI progressively increases from level walking to ramp descent and stair descent, a pattern that is consistently reproduced by all four muscle redundancy solvers.

To assess the relationship between KCF errors and muscle co-contraction, we further analyzed the correlation between RMSE and CCI using a linear regression model with cluster-robust standard errors in Figure 5. This analysis was motivated by previous results showing that CCI-informed simulations have lower RMSE values than the minimum activation (RMR) solution, particularly in subjects with higher co-contraction. The corresponding regression coefficient (*β*, in BW per unit CCI), cluster-robust standard errors, 95 % confidence intervals (95 % CI), *p*-values, and the number of observations (*N*) are reported in Table S4 of the Supplementary Material. Furthermore, the results of the linear mixed-effect model for all converged analyses are reported in Table S5 of the Supplementary Material. The results in Figure 5a indicate a positive correlation between RMSE and CCI for the minimum activation (RMR) simulations (*β* = 1.55, 95% CI [0.79, 2.30]), whereas the correlation in the CCI-informed muscle redundancy solver is smaller (*β* = 0.78, 95% CI [0.14, 1.43]). In Figure 5b-d, the correlation between

**Figure 5.**
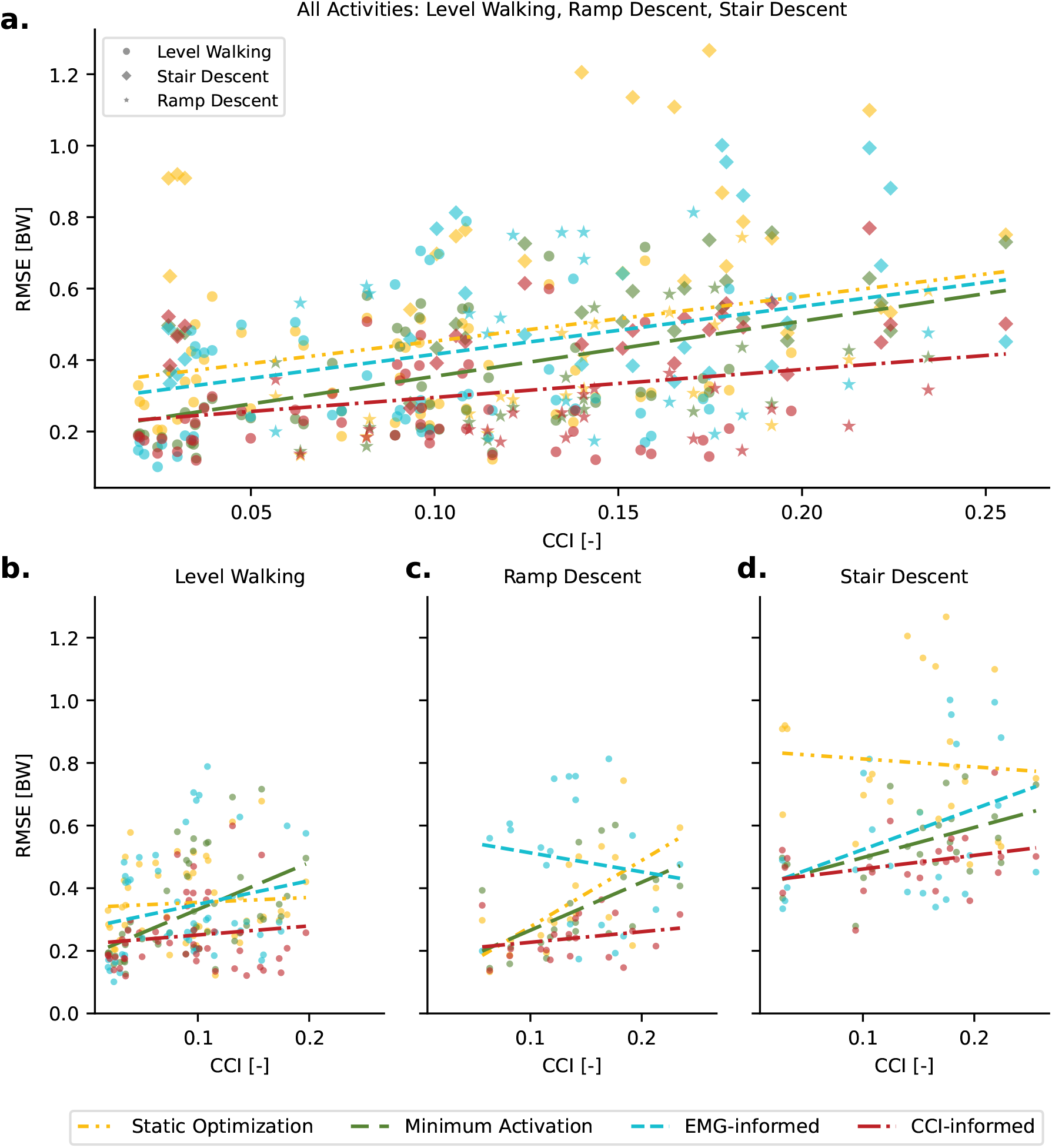
Statistical analysis of the correlation between RMSE and CCI using a linear regression with cluster-robust standard errors, comparing the four muscle redundancy solvers across all activities combined (a) and for each activity separately (b–d). A positive association (*β* = 1.55, 95% CI [0.79, 2.30]) between RMSE and CCI was observed for the minimum activation (RMR) solver across all activities, as well as for each activity individually (*β >* 0.96). In contrast, this association was weaker for the CCI-informed solver across all activities (*β* = 0.78, 95% CI [0.14, 1.43]) and specifically during level walking and ramp descent, with *β* = 0.29 (95% CI [-1.34, 1.92]) and *β* = 0.34 (95% CI [-0.50, 1.17]), respectively.

RMSE and CCI is plotted separately for each activity. These results suggest that the CCI-informed method, particularly for level walking (*β* = 0.29, 95% CI [-1.34, 1.92]) and ramp descent (*β* = 0.34, 95% CI [-0.50, 1.17]), largely eliminates the correlation between RMSE and CCI observed with the minimum activation (RMR) solver. The linear mixed-effects results indicated broadly consistent trends between the CCI-informed and minimum activation (RMR) solvers for converged analyses, with the minimum activation (RMR) solver generally showing positive associations between CCI and RMSE, whereas the CCI-informed solver exhibited weaker effects. However, accounting for between-subject variability in the mixed-effects models generally led to larger effect estimates.

## Discussion

The aim of this study was to evaluate the performance of four different muscle redundancy solvers for estimating KCF from motion capture measurements. We tested their estimates against *in vivo* joint force measurements obtained from an instrumented implant. Specifically, we included two commonly used muscle redundancy solvers, SO and RMR, which both minimize muscle activations. The key difference between these solvers is that passive muscle forces are incorporated in the RMR solver but not in the SO solver. To account for changes in muscle coordination strategies, we additionally proposed two EMG-based approaches. The EMG-informed approach minimizes the error between estimated and measured muscle activations. The CCI-informed approach minimizes the error in the level of co-contraction between the VL and MG muscle pair, thereby reducing the experimental setup required to account for changes in muscle coordination strategies.

In this study, we used a fixed set of weights in the objective function of the RMR solver. The EMG- and CCI-specific terms in the objective function were assigned weights twice as large as those of the muscle activation minimization term to account for variations in muscle coordination strategies. Reserve actuators were strongly penalized by assigning weights 10 times higher than those of the muscle activation minimization term to ensure a minimal contribution of reserve actuators. For both EMG-informed and CCI-informed simulations, weighting factors were not tuned to improve agreement with measured *in vivo* knee joint contact forces, to avoid biasing the results toward the measured knee joint contact forces, which are not available in practice, and to preserve the generalizability of the proposed approach.

Our results indicate the CCI-informed simulations performed most consistently across subjects, resulting in the best performance overall across all three evaluation metrics (RMSE, *R*^2^, and peak force error) for all three activities with the fixed set of weights in the objective function. Approximately 50–75% of the compressive KCF is contributed by tensile muscle forces acting across the joint^9^, indicating that changes in muscle coordination strategies can substantially impact KCF^36^. By minimizing the CCI error in the objective function of the muscle redundancy solver, subject-specific coordination patterns of VL and MG are incorporated into the KCF estimate. This is reflected in the RMSE results for subjects K1L, K2L, K5R, and K7L, where a larger discrepancy between measured CCI and CCI estimated from minimum activation (RMR) simulations was observed. For these subjects, the CCI-informed approach resulted in significantly lower RMSE (*p <* 0.01) and higher *R*^2^ values compared to the minimum activation (RMR) solver. These results are further supported by the correlation analysis between RMSE and CCI in Figure 5. In the combined plot across all activities (Figure 5a), the results indicate a positive correlation between CCI and RMSE for the minimum activation (RMR) simulations, whereas the correlation is reduced in the CCI-informed simulations. The effect of co-contraction becomes even more pronounced when each activity (Figure 5b-d) is analyzed separately, because median co-contraction levels increase progressively from level walking to ramp descent and stair descent in the EMG measurements (Figure 4b). Overall, the CCI-informed simulations effectively capture co-contraction variations, resulting in more consistent RMSE values across different co-contraction levels without further tuning of the cost function weights.

Furthermore, when comparing the two EMG-based approaches, the EMG-informed simulations showed greater variability across subjects compared to the CCI-informed approach: for example, RMSE values were lowest for three subjects but highest for the remaining three, and we attribute the high variability to inaccurate MVC normalization, potentially due to muscle inhibition during MVC measurements^37^. Consequently, muscle activations in EMG-informed cases can be easily overestimated, leading to elevated KCF. In contrast, the CCI-informed approach is less sensitive to the absolute magnitude of the MVC used for EMG normalization than the EMG-informed solver, because the CCI is defined as a ratio of muscle activations. If MVC values are underestimated similarly across the muscle pair, the relative activation between the muscles remains largely preserved. Consequently, the CCI approach can be more robust than EMG-informed simulations that rely on absolute EMG amplitudes.

Comparison of the two minimum activation approaches, SO and RMR, revealed that SO overpredicted the second KCF peak for subject K8L during level walking and for multiple subjects during ramp and stair descent (Figure 2). Whittington et al. previously reported that approximately 35% of the net hip flexor moment during push-off in level walking arises from passive hip flexor stretch^38^. Passive contributions from the ankle plantar flexors are around 10%^38^. When these passive forces are not accounted for, additional muscle activation is required, including biarticular muscles crossing the knee, thereby increasing muscle co-contraction and the resulting KCF, with the elevated co-contraction reflecting model limitations rather than actual muscle behavior.

In addition, our results indicate that higher RMSE and peak force errors across all muscle redundancy solvers are estimated during stair descent compared to level walking and ramp descent (Figure 4 and Supplementary Material Table S2). One possible reason is that the prescribed secondary knee motions, derived from passive flexion measurements in cadavers^39^, may not be accurately captured during activities involving greater knee flexion, such as stair descent. A previous study using a 6-DOF knee model to simulate squatting from the CAMS dataset showed substantial improvement in the KCF RMSE compared to a study that used a 1-DOF model^40,41^. Furthermore, Nejad et al. reported an overprediction of KCF, particularly in deep knee and hip flexion^41^. These results emphasize the role of activity-specific knee kinematics and motivate the development of knee models that more comprehensively represent multiplane joint motion across different activities, while not being as computationally demanding as current complex 6-DOF knee joint models^42^.

Our estimated KCF and corresponding RMSE values are similar to those reported in previous studies of the CAMS dataset. Pellegrinelli et al.^43^ reported RMSE values of 0.37 BW for level walking using SO, which is very close to our results of 0.35 BW during level walking. Slightly higher RMSE values of 0.44 BW and 0.48 BW in level walking have been reported by Guitteny et al.^44^ and Nejad et al.^41^, respectively. This difference may be attributed to differences in the underlying MSK models used in both studies. Furthermore, Guo et al.^40^ developed a subject-specific model for one participant from the CAMS dataset and reported strong agreement with *in vivo* measurements, with RMSE values below 0.16 BW for level walking and squatting. This high level of accuracy is likely attributable to subject-specific alignment and bone and implant positioning using computed tomography scans. In addition, the knee joint was modeled with 6 DOF at the tibiofemoral joint, and flexion–extension was prescribed based on fluoroscopy measurements. Together, these approaches enabled a more accurate representation of implant-specific kinematics. The knee kinematic improvements influence both muscle force and contact force predictions by altering muscle moment arms and the center of pressure location relative to the joint center, which in turn affects the muscle forces required to satisfy dynamic equilibrium and the resulting estimated KCF. As inaccuracies in joint kinematics have been shown to translate into errors in the estimated KCF, accounting for subject-specific anatomical and kinematic characteristics likely contributed to the high agreement with *in vivo* measurements^45^.

Compared to the previously developed EMG-assisted approach applied to the Knee Grand Challenge dataset^46^, the improvements observed in this study are of similar magnitude. Bennett et al.^23^ and Esrafilian et al.^47^ reported mean RMSE reductions of approximately 0.11 BW and 0.09 BW, respectively, when comparing SO with the EMG-assisted approach using a one-DOF knee model. This is comparable to the 0.10 BW improvement achieved in the present study. In contrast, Princelle et al. reported substantially smaller reductions, with mean RMSE improvements below 0.04 BW^48^. Notably, unlike the EMG-assisted approach, the CCI-informed approach does not require model calibration, a process that has been reported to be time-intensive^23^. Thus, comparable improvements are achieved at a substantially lower computational cost and greater ease of use, while relying on fewer EMG input measurements.

The present study has several limitations. First, the sample size was limited to six participants from the CAMS dataset. The availability of datasets with instrumented implants is limited by ethical and practical constraints, hindering validation across larger, more diverse populations. Nevertheless, the CAMS dataset remains one of the most comprehensive resources, providing multiple activities alongside motion capture, ground reaction forces, EMG, and fluoroscopy measurements. In addition, statistical inference was limited by the small number of data clusters, which may affect the reliability of standard error estimates, and results are therefore interpreted primarily in terms of effect direction and magnitude rather than formal statistical significance. Future simulations and comparisons using additional datasets, such as the Knee Grand Challenge dataset^46^, would further validate the approach.

Second, our analysis was limited to level walking, ramp descent, and stair descent, while activities such as sit-to-stand and squatting were excluded. Previous studies have shown that performing the simulations with the one-DOF Walker knee joint tends to overpredict knee contact forces at deep knee and hip flexion angles, regardless of the changes in muscle coordination strategies^39,41^. This is likely due to limitations in the prescribed knee kinematics, as accurate kinematic representation has been shown to improve agreement with estimated knee joint forces^40^. Further development of knee joint models that better capture natural internal rotation and abduction motion may help achieve similar accuracy in KCF estimation for activities involving high knee and hip flexion angles. Furthermore, this study focused solely on validating compressive KCF. In many weight-bearing tasks, compressive knee joint forces are larger than shear forces and are therefore the main contributor to overall knee joint loading^26^. However, additional validation of other force components is also necessary, as these may provide further insight into changes in knee joint loading associated with KOA.

Finally, the CCI-informed approach targeted only the co-contraction between the VL and MG, as these muscles showed the largest discrepancies between measured and minimum-activation–predicted co-contraction in the studied subjects. Consequently, changes in coordination strategies of other muscle groups are only captured to a limited extent. Further investigation is required to determine whether similar patterns are observed in other patient populations, such as individuals with early-stage KOA.

In conclusion, our results demonstrate that the CCI-informed approach consistently achieves lower RMSE values for estimating KCF across all activities and subjects compared to the minimum activation and full EMG-informed methods. Furthermore, the CCI-informed approach captures changes in muscle coordination strategies that directly impact estimated KCF. Changes in coordination were often not detected by minimizing muscle activation alone, leading to higher RMSE values at higher co-contraction levels. In addition, the CCI-informed method is more feasible to apply, as it requires EMG measurements from only two muscles to account for changes in muscle coordination strategies, which could be measured in functional clothing. This makes it particularly promising for applications outside the laboratory to track rehabilitation methods, including gait retraining.

## Acknowledgements

This publication is part of the project LoaD (projectnr. NWA1389.20.009) of the NWA-ORC research programme which is (partly) financed by the Dutch Research Council (NWO).

## Author contributions statement

S.H. and A.S. conceptualized the study. S.H. performed the MSK simulations, analyzed the results, and created the illustrations and figures. All authors aided in data interpretation. S.H. drafted the manuscript, and all authors critically reviewed and approved the final manuscript.

## Code availability

The Rapid Muscle Redundancy solver code including the EMG-informed and CCI-informed approach is available at https://github.com/ComputationalBiomechanicsLab/rmr-solver-py.git.

## Data availability

The CAMS dataset is publicly available and can be accessed upon request via the OrthoLoad website: https://cams-knee.orthoload.com/data/data-request/

## Additional information

Supplementary Information. The online version contains supplementary material.

