## Supplementary Material Table S1-S5 for "Co-contraction index improves estimates of knee joint contact forces: A musculoskeletal modelling study"

| Subject | SO vs CCI | SO vs EMG | SO vs RMR | CCI vs EMG | CCI vs RMR | EMG vs RMR |
|---|---|---|---|---|---|---|
| K1L | $< 0.01$ | $< 0.01$ | $< 0.01$ | $< 0.01$ | $< 0.01$ | $< 0.01$ |
| K2L | $< 0.01$ | 0.29 | 0.29 | $< 0.01$ | $< 0.01$ | 0.20 |
| K3R | $< 0.01$ | $< 0.01$ | $< 0.01$ | $< 0.01$ | 0.73 | $< 0.01$ |
| K5R | $< 0.01$ | $< 0.01$ | 0.81 | 0.86 | $< 0.01$ | $< 0.01$ |
| K7L | $< 0.01$ | $< 0.01$ | 0.04 | $< 0.01$ | 0.04 | $< 0.01$ |
| K8L | $< 0.01$ | 0.29 | $< 0.01$ | $< 0.01$ | 0.50 | $< 0.01$ |
| Overall | $< 0.01$ | 0.23 | $< 0.01$ | $< 0.01$ | $< 0.01$ | 0.15 |

**Table S1. Pairwise *p*-values comparing KCF estimation errors (RMSE) across four muscle redundancy solvers, by subject and across all subjects combined (Overall).** The evaluated solvers were the Static Optimization (SO), original Residual Muscle Redundancy (RMR), EMG-Informed (EMG), and CCI-Informed (CCI) solver. The *p*-values are reported from Wilcoxon signed-rank tests with Holm-Bonferroni correction for multiple comparisons. The *p*-values highlighted in green indicate statistically significant differences in RMSE between compared solver pairs ($p < 0.05$).

| Metric | Activity | Static Optimization | Minimum Activation | EMG-informed | CCI-informed |
|---|---|---|---|---|---|
| RMSE [BW] (mean ± std) | Level Walking | $0.35 \pm 0.12$ | $0.32 \pm 0.15$ | $0.34 \pm 0.18$ | $0.25 \pm 0.11$ |
| | Ramp Descent | $0.36 \pm 0.16$ | $0.33 \pm 0.14$ | $0.49 \pm 0.21$ | $0.24 \pm 0.07$ |
| | Stair Descent | $0.80 \pm 0.23$ | $0.54 \pm 0.12$ | $0.58 \pm 0.23$ | $0.48 \pm 0.10$ |
| | Overall | $0.47 \pm 0.25$ | $0.38 \pm 0.17$ | $0.44 \pm 0.22$ | $0.31 \pm 0.14$ |
| $R^2$ [-] (mean ± std) | Level Walking | $0.84 \pm 0.10$ | $0.88 \pm 0.08$ | $0.84 \pm 0.14$ | $0.93 \pm 0.05$ |
| | Ramp Descent | $0.89 \pm 0.08$ | $0.91 \pm 0.08$ | $0.78 \pm 0.15$ | $0.95 \pm 0.03$ |
| | Stair Descent | $0.45 \pm 0.34$ | $0.77 \pm 0.08$ | $0.71 \pm 0.19$ | $0.81 \pm 0.09$ |
| | Overall | $0.75 \pm 0.26$ | $0.86 \pm 0.10$ | $0.80 \pm 0.17$ | $0.90 \pm 0.08$ |
| Peak Force Error [BW] (mean ± std) | Level Walking | $0.42 \pm 0.42$ | $0.40 \pm 0.30$ | $0.36 \pm 0.27$ | $0.33 \pm 0.27$ |
| | Ramp Descent | $0.63 \pm 0.54$ | $0.34 \pm 0.24$ | $0.50 \pm 0.32$ | $0.28 \pm 0.18$ |
| | Stair Descent | $2.06 \pm 0.85$ | $0.71 \pm 0.41$ | $0.74 \pm 0.52$ | $0.64 \pm 0.37$ |
| | Overall | $0.89 \pm 0.91$ | $0.47 \pm 0.35$ | $0.49 \pm 0.39$ | $0.40 \pm 0.32$ |

**Table S2. Summary of RMSE in body weight (BW), $R^2$ and peak force error of the second peak in stance phase for each activity and overall across all activities.** Metrics are compared across four muscle redundancy solvers: Static Optimization, Minimum Activation (RMR), EMG-informed, and CCI-informed. For each activity and the overall results, the best-performing solver is highlighted in green and the worst-performing solver in red. The results show that the CCI-informed simulations performed overall the best in each evaluation metric.

| Activity | SO vs CCI | SO vs EMG | SO vs RMR | CCI vs EMG | CCI vs RMR | EMG vs RMR |
|---|---|---|---|---|---|---|
| Level Walking | $< 0.01$ | 0.36 | 0.37 | 0.08 | $< 0.01$ | 0.85 |
| Ramp Descent | $< 0.01$ | 0.14 | 0.37 | $< 0.01$ | $< 0.01$ | 0.08 |
| Stair Descent | $< 0.01$ | 0.03 | $< 0.01$ | 0.10 | $< 0.01$ | 0.55 |

**Table S3. Pairwise *p*-values comparing KCF estimation errors (RMSE) across four muscle redundancy solvers by activity.** The evaluated solvers were the Static Optimization, original Residual Muscle Redundancy (RMR), EMG-Informed (EMG), and CCI-Informed (CCI) solver. The *p*-values are reported from Wilcoxon signed-rank tests with Holm-Bonferroni correction for multiple comparisons. The *p*-values highlighted in green indicate statistically significant differences in RMSE between compared solver pairs ($p < 0.05$).

| Activity | Solver | $\beta$ | Standard Error | 95% CI | $p$-value | $N$ (observations) |
|---|---|---|---|---|---|---|
| All Activities | Static Optimization | 1.25 | 0.70 | [-0.55, 3.06] | 0.13 | 92 |
| | Minimum Acitvation | 1.55 | 0.30 | [0.79, 2.30] | 0.00 | 92 |
| | EMG-Informed | 1.34 | 0.76 | [-0.60, 3.28] | 0.14 | 92 |
| | CCI-Informed | 0.78 | 0.25 | [0.14, 1.43] | 0.03 | 92 |
| Level Walking | Static Optimization | 0.16 | 0.69 | [-1.62, 1.94] | 0.83 | 48 |
| | Minimum Acitvation | 1.50 | 0.64 | [-0.14, 3.14] | 0.07 | 48 |
| | EMG-Informed | 0.75 | 1.22 | [-2.39, 3.90] | 0.56 | 48 |
| | CCI-Informed | 0.29 | 0.63 | [-1.34, 1.92] | 0.67 | 48 |
| Ramp Descent | Static Optimization | 2.12 | 0.45 | [0.88, 3.36] | 0.01 | 20 |
| | Minimum Acitvation | 1.53 | 0.35 | [0.56, 2.51] | 0.01 | 20 |
| | EMG-Informed | -0.61 | 0.62 | [-2.33, 1.12] | 0.38 | 20 |
| | CCI-Informed | 0.34 | 0.30 | [-0.50, 1.17] | 0.32 | 20 |
| Stair Descent | Static Optimization | -0.25 | 0.75 | [-2.19, 1.68] | 0.75 | 24 |
| | Minimum Acitvation | 0.96 | 0.36 | [0.03, 1.89] | 0.04 | 24 |
| | EMG-Informed | 1.30 | 0.88 | [-0.96, 3.55] | 0.20 | 24 |
| | CCI-Informed | 0.43 | 0.37 | [-0.53, 1.39] | 0.30 | 24 |

**Table S4. Results of linear regression analyses with cluster-robust standard errors examining the association between the CCI and the RMSE of estimated compressive KCF for each solver and activity condition.** The six subjects were treated as clusters. Regression coefficients ($\beta$, in BW per unit CCI), cluster-robust standard errors, 95 % confidence intervals (95 % CI), $p$-values, and the number of observations ($N$) are reported for the Static Optimization, Minimum Activation (RMR), EMG-informed, and CCI-informed solvers across all activities combined, as well as separately for level walking, ramp descent, and stair descent.

| Activity | Solver | $\beta$ | RE SD | 95% CI | $R^2_{\text{marg}}$ | ICC | $N$ (observations) |
|---|---|---|---|---|---|---|---|
| All Activities | Static Optimization | 2.44 | 0.14 | [0.96, 3.66] | 0.23 | 0.30 | 92 |
| | Minimum Acitvation | 1.94 | 0.11 | [1.55, 2.73] | 0.37 | 0.52 | 92 |
| | EMG-Informed | 1.47 | 0.18 | [1.03, 2.31] | 0.14 | 0.69 | 92 |
| | CCI-Informed | NA$^{\dagger}$ | NA | NA | NA | NA | NA |
| Level Walking | Static Optimization | 1.83 | 0.14 | [0.82, 2.88] | 0.24 | 0.78 | 48 |
| | Minimum Acitvation | 2.62 | 0.14 | [1.66, 3.24] | 0.42 | 0.83 | 48 |
| | EMG-Informed | -0.31 | 0.19 | [-0.83, 1.05] | 0.01 | 0.94 | 48 |
| | CCI-Informed | 0.90 | 0.11 | [0.01, 1.64] | 0.11 | 0.75 | 48 |
| Ramp Descent | Static Optimization | NA$^{\dagger}$ | NA | NA | NA | NA | NA |
| | Minimum Acitvation | 1.18 | 0.12 | [0.64, 1.77] | 0.16 | 0.93 | 20 |
| | EMG-Informed | 0.71 | 0.22 | [0.23, 1.69] | 0.02 | 0.91 | 20 |
| | CCI-Informed | NA$^{\dagger}$ | NA | NA | NA | NA | NA |
| Stair Descent | Static Optimization | 0.60 | 0.22 | [-1.00, 3.67] | 0.02 | 0.80 | 24 |
| | Minimum Acitvation | NA$^{\dagger}$ | NA | NA | NA | NA | NA |
| | EMG-Informed | 0.77 | 0.21 | [-0.20, 2.12] | 0.04 | 0.80 | 24 |
| | CCI-Informed | NA$^{\dagger}$ | NA | NA | NA | NA | NA |

**Table S5. Results of the linear mixed-effects regression analysis between the CCI and the RMSE of estimated compressive KCF for each solver and activity condition.** The fixed-effect slope ($\beta$, in BW per unit CCI), its 95 % confidence interval (95 % CI), the between-subject random-effect standard deviation (RE SD), the marginal coefficient of determination ($R^2_{\text{marg}}$, variance explained by CCI alone), the intraclass correlation coefficient (ICC, proportion of total variance attributable to between-subject differences), and the number of observations ($N$) are reported for Static Optimization, Minimum Activation (RMR), EMG-informed, and CCI-informed solvers across all activities combined, as well as separately for level walking, ramp descent, and stair descent. The mixed-effects model included subject as a random intercept to account for between-subject variability in baseline RMSE across repeated trials. Models that did not converge are indicated by $^{\dagger}$ and were excluded from the analysis.
